# Open or closed? Exploring the conformational heterogeneity of human plasminogen across multiple resolution scales

**DOI:** 10.64898/2026.07.23.740275

**Authors:** Linda Kasiarova, Naina Verma, Matyas Pinkas, Tomas Henek, Tomas Klumpler, Marie Chovancikova, Helena Reutova, Petr Kasparek, Martina Hlozankova, Jiri Damborsky, Jiri Novacek, Lenka Hernychova, Martin Marek

**Affiliations:** Loschmidt Laboratories, Department of Experimental Biology and RECETOX, Faculty of Science, Masaryk University, Kamenice 5, Bld. C13, 625 00 Brno, Czech Republic; International Clinical Research Center, St. Anne’s University Hospital Brno, Pekarska 53, 656 91 Brno, Czech Republic; Research Center for Applied Molecular Oncology, Masaryk Memorial Cancer Institute, Zluty kopec 7, 656 53 Brno, Czech Republic; Central European Institute of Technology, Masaryk University, Kamenice 753/5, 625 00 Brno, Czech Republic; BioVendor LM, Karasek 1767/1, 621 00 Brno, Czech Republic

**Keywords:** plasminogen, fibrinolysis, staphylokinase, conformational landscape, cryo-EM, HDX-MS, SAXS

## Abstract

Plasminogen (Plg), the zymogen of the fibrinolytic protease plasmin, is a multidomain, conformationally-rich protein that plays a crucial role in targeted thrombolysis for stroke and heart attack treatments. Yet, the conformational state that affects binding to Plg activators and fibrinolytic activity is rarely validated, contributing to poor reproducibility and failures in drug development. Here, we establish a structural biology workflow to characterize human Plg suspended in an aqueous solution, preserving it in a near-native state and eliminating the need for crystal growth. Two variants, cleavage-resistant Plg-RV (R561A) and catalytically inactive Plg-CAT (S741A), were expressed in mammalian cells. Both variants were correctly folded and stable (T_m_ ≈ 60.6°C) and, unlike commercial plasma-derived Plg, were resistant to staphylokinase-mediated activation. Small-angle X-ray scattering and hydrogen-deuterium exchange mass spectrometry revealed substantial conformational heterogeneity. The recombinantly-produced Plg variants adopted a closed conformation, exhibiting a good fit to the closed structure determined by X-ray diffraction. Conversely, the plasma-derived Plg populated an extended, open-like state. Cryo-EM analysis of the Plg-RV variant yielded a 4.4 Å resolution map, and a rigid-body-fitted model revealed the closed-state architecture. Our findings demonstrate that rigorous structural validation of Plg is essential for future functional studies and rational development of next-generation thrombolytic agents.

## Introduction

Human plasminogen (Plg) is the inactive zymogen of plasmin, a central protease of the fibrinolytic system responsible for clot dissolution and a wide spectrum of physiological and pathological processes^1,2^. In addition to its role in endogenous fibrinolysis, Plg is exploited by several bacterial pathogens, including *Staphylococcus aureus*, which hijack the host fibrinolytic machinery to promote tissue invasion. *S*. *aureus* produces staphylokinase (SAK), a bacterial plasminogen activator, that forms a functional complex with plasmin (Plm) and Plg, leading to efficient conversion of Plg into an active Plm^3,4^.

Structurally, Plg is a multidomain protein composed of an N-terminal Pan-apple (PAp) domain, five kringle domains, and a C-terminal serine protease domain^5^. The relative arrangement of these domains gives rise to distinct global conformational states, commonly referred to as closed and open conformations^5,6^, which distinctly impact activation, ligand binding, and complex formation^7^. The crystal structures of uncomplexed full-length human Plg showed a compact and closed conformation of the Plg zymogen^5^.

Despite the central role of Plg activation in SAK-mediated fibrinolysis, structural knowledge on the complexation between the full-length Plg and SAK is essentially missing. Structural probing of the Plg-SAK complexation is challenging due to the high conformational flexibility of multi-domain Plg and the transient nature of the complex formation^8^. Importantly, Plg exists in an equilibrium between the compactly closed and the extended open conformations, which differ substantially in ligand accessibility and activation propensity^5–7^. Despite this fact, the conformational states of Plg preparations are rarely experimentally validated before structural and biochemical experiments. This may contribute to inconsistencies in biological conclusions and low reproducibility of results across laboratories that use Plg specimens of various origins.

So far, only one co-crystal structure of the SAK-bound Plm complex is available^8^. However, it only captures a ternary complex of SAK-bound microplasmin (mPlm), a truncated construct comprising only the catalytic serine protease domain, lacking the Pan-apple domain and kringle domains^8^. While this structure provided some mechanistic insights into SAK-mediated Plm activation, it does not reflect the behavior of the full-length Plg-SAK complex. Recently, small-angle X-ray scattering (SAXS) technique has been employed to probe the conformational state of Plg, highlighting the key role of biophysical quality control before downstream structural and biochemical studies^6^.

Recent advances in cryogenic electron microscopy (cryo-EM) have enabled structural characterization of increasingly smaller and more flexible proteins under near-native, crystal-lattice-free conditions^9,10^. Establishing structurally validated and conformationally homogeneous plasminogen preparations is therefore a critical prerequisite for meaningful investigation of the Plg-SAK system. While SAXS and cryo-EM report on the global architecture of Plg, they do not resolve the local backbone dynamics that distinguish the activation-competent open state from the autoinhibited closed state, nor they identify the local structural regions affected by SAK interaction and activation. To access this information, we employed hydrogen-deuterium exchange mass spectrometry (HDX-MS), which probes solvent accessibility and conformational flexibility at peptide-level resolution^11^.

In this work, we establish an integrative structural biology approach, ranging from mammalian recombinant production and biochemistry characterisation to structural exploration (SAXS, cryo-EM, and HDX-MS), for mechanistic studies of Plg. Importantly, we demonstrate that recombinantly produced full-length Plg variants consistently adopt a compact closed conformation that is distinct from those found in commercially available plasma serum-sourced Plg. We utilized HDX-MS to correlate global conformational states with local protection patterns in the activation-loop region and to characterize SAK-associated structural changes. Crucially, we demonstrate that the conformational homogeneity determines suitability for single-particle cryo-EM analysis. Our results thus establish a framework for structural and mechanistic studies of SAK-based Plg activation.

## Materials and methods

### Design of Plg variants

Human Plg R561A (Plg-RV) and Plg S741A (Plg-CAT) were designed for specific functional purposes. The Plg-RV variant contains a single point mutation at the cleavage site required for activation of Plg to Plm. This mutation was introduced to prevent proteolytic cleavage, thereby maintaining Plg in its inactive zymogen form even in the presence of activation factors such as SAK. The Plg-CAT variant was designed to carry a single point mutation within the catalytic triad (His603, Asp646, Ser741), specifically targeting the active-site serine residue (Ser741). In both variants, the selected amino acid residues were substituted with alanine to abolish their functional contribution. A schematic representation of the designed variants is shown in **Fig. S1**.

### Gene synthesis and molecular cloning

Plg-RV and Plg-CAT were assembled by *de novo* synthesis (GenScript Biotech) into plasmid pUC57, streamlined for transcription and adapted to mammalian codon usage. Gene synthesis included the addition of flanking restriction sites for KpnI and XhoI required for subcloning into the expression vector pcDNA3.1. The constructs were designed to carry a C-terminal 6xHis tag and N-terminal signal peptide (MWWRLWWLLLLLLLLWPMVWA). Plasmids were isolated using the MEGA prep kit (Qiagen) for small-scale transfections and using the GIGA prep kit (Qiagen) for large-scale transfections. SAK-Star-WT gene was assembled as previously reported^12^.

### Protein overproduction and purification

Protein production was carried out via transient co-transfection of the expression vector pcDNA3.1-Plg-RV/pcDNA3.1-Plg-CAT, together with a helper plasmid and pmaxGFP as a transfection efficiency marker. Suspension-adapted human embryonic kidney (HEK293) cell line that stably expresses the Epstein-Barr virus nuclear antigen 1 (EBNA1) was used for protein production.

Before scaling up, transfection conditions were optimized in a small-scale format using a 12-well plate. On the day of transfection, the required number of cells was harvested by centrifugation at 250 × g, 24 °C, 10 min, and the pellet was resuspended in Gibco® FreeStyle™ 293 Expression Medium at a final concentration of 2 × 10 cells/mL. A volume of 500 μL cell suspension was added per well. Plasmid DNA and polyethylenimine (PEI) were added to each well, with PEI added last. After DNA addition, the cells were incubated at 198 rpm, 37 °C, in an incubator supplemented with CO for 4 hours. Following this incubation, cells were diluted 1:12 with HyClone medium supplemented with 2 mM L-glutamine, and 0.3 mM sodium butyrate, and returned to the incubator under shaking conditions (198 rpm, 37 °C). Over the next several days, cell density, viability, and GFP expression were monitored microscopically. On day 5, the cultivation was terminated and efficiency and quality of the transfection were assessed.

Based on the results of the small-scale optimization, large-scale production was carried out under similar conditions in 2-4 L cultivation medium. The large-scale transfection protocol mirrored the small-scale setup, including media composition, PEI-mediated transfection, and post-transfection handling. Cultivation was performed in glass square bottles at 110 rpm and 37 °C. At the end of cultivation, cells were harvested by centrifugation at 4500 × g, 4 °C, 20 min. The resulting supernatant, containing the secreted Plg variant, was directly subjected to purification.

Before purification, the culture medium containing the secreted protein was filtered (1.6-0.6 µm) and concentrated using a Vivaflow 200 Cross Flow Cassette, to a volume of 50-200 mL, depending on the initial volume of the culture media. The concentrated medium was then loaded onto a 5 mL Ni^2^ - NTA affinity chromatography column (Macherey-Nagel). Elution of plasminogen was observed at an imidazole concentration of 24% as a gradient of buffer B. The purification process used two buffers: buffer A (50 mM Tris, 300 mM NaCl, pH = 8) and buffer B (50mM Tris, 300 mM NaCl and 250mM imidazole, pH = 8.0). The purified protein was then dialyzed against 10 L of dialysis buffer (20 mM Tris and 20mM NaCl, pH = 8). The dialysis buffer was exchanged after 24 hours, with the full dialysis lasting 48 hours. The eluate from affinity purification was then subjected to gel filtration chromatography. Recombinant SAK-Star-WT was produced and purified as previously described^12^, and its quality was assessed as previously described^12^. The recombinant expression system used for Plg variants production is proprietary to BioVendor and is available under a material transfer agreement.

### Western blotting

Western blotting (WB) was performed to verify whether the target protein was successfully produced. For detection, an anti-His antibody (Qiagen cat. no. 34670, 1:1000) and an HRP-conjugated anti-mouse secondary antibody (Agilent P044701-2, 1:1000) were used. To specifically monitor Plg production, an anti-Plg antibody (ab154560, 1:1000) and an HRP-conjugate anti-rat secondary antibody were used (Agilent P044801-2, 1:1000). Anti-Plg antibody was specifically used to determine whether the Plg-CAT contains fragments corresponding to plasmin (Plg to Plm activation happens through the cleavage step).

### Biophysical protein quality controls

Purity of proteins was checked by 10% acrylamide Tris-glycine SDS-PAGE. Concentration was determined by absorbance at 280nm, assuming molar extinction coefficient of 152 200 M-1.cm-1 for both Plg-RV and Plg-CAT (Expasy, ProtParam). Protein thermal stability was assessed using NanoDifferential Scanning Fluorimetry (nanoDSF) on a Prometheus instrument at 1°C/min. Melting temperatures (Tm) were determined using the first derivative of the 350/330 nm fluorescence ratio curve. CD spectra were obtained at 0.2 mg/mL final concentration of Plg-RV and Plg-CAT from 195 to 280 nm at 1nm bandwidth with 0.5 s integration time as an average of 3 readings using a Chirascan spectropolarimeter (Applied Photophysics, UK). Content of secondary structural elements was assessed from CD spectra using BeStSel software^13^.

### SAXS data collection

For SAXS measurement Plg-RV, Plg-CAT and Plg-WT were concentrated to 2.5 mg/mL in case of Plg-WT and 5 mg/mL for Plg-RV and Plg-CAT and dialyzed into PBS buffer (10mM Na_2_HPO_4_, 1.8 mM KH_2_PO_4_, 2.7 mM KCl, 137 mM NaCl, pH 7.4). SAXS data was collected using a Rigaku BioSAXS-2000 instrument at CEITEC (Brno, Czech Republic) equipped with a HyPix-3000 detector at a sample-detector distance of 0.5 m in q range 0.009-0.65 Å^-1^ (I(s) vs q, where q = 4π sinθ/λ; 2θ is the scattering angle and λ = 1.54 Å). Six 10 min frames were collected at a sample temperature of 20°C. The datasets were normalized to the intensity of the transmitted beam and radially averaged. Scattering curves from individual frames were checked for radiation damage and averaged. The corresponding scattering from the solvent-blank was subtracted to produce the scattering profile.

### SAXS data analysis and deposition

Subtracted SAXS datasets were processed using the ATSAS 2.8.3 software package^14^. Integral structural parameters determined by PRIMUS^15^ are summarized in **Table S1**. Evaluation of atomic models and fitting to experimental data was performed by CRYSOL^16^. *Ab initio* reconstruction was done using DAMMIN^17^, where the computation mode was set to slow, while all other parameters were kept at their default values. The SAXS data were deposited at SASBDB^18^ under the accession codes: SASDZU3, SASDZ24, SASDZ34.

### Sample preparation for cryo-EM

The overall volume of 3.5 μL of purified Plg-RV (0.47 mg/mL) was mixed with SAK in 1:2 (1 Plg : 2 SAK) molar ratio, and sample was applied to a freshly plasma-cleaned (60s H/O plasma, Ametek Solarus II) Quantifoil R1.2/R1.3 300-mesh copper grids (Quantifoil Micro Tools GmbH, Jena, Germany). After 15 s incubation period, the grids were plunge-frozen in liquid ethane using the Vitrobot Mark IV (Thermo Fisher Scientific) vitrification device kept at 4°C and 100% humidity.

### Cryo-EM data collection

The TEM grids were first screened in a Talos Arctica (Thermo Fisher Scientific) electron microscope, operated at 200 kV and equipped with the Falcon IV direct electron detector and the Selectris energy filter (Thermo Fisher Scientific). A small data set was collected to evaluate sample quality before transferring the selected grids to a 300 kV Titan Krios (Thermo Fisher Scientific) transmission electron microscope. The microscope was aligned for parallel and fringe-free illumination. A data set consisting of 14 330 movies was collected on a K3 direct electron detector with the Bioquantum energy filter (Gatan, Inc.) with the slit width set to 10 eV, using the SerialEM software^19^. Each movie consisted of 40 frames with a frame dose of 1.4 e /A^2^ and the pixel size of 0.5113 A/px. The nominal defocus range was set to −1.0 to −2.4 μm. All data collection parameters are summarized in **Table S2**.

### Processing and 3D mapping of cryo-EM images

Data processing was carried out using a standard pipeline in CryoSPARC^20^ (**Fig. S2**). Movies were aligned using Patch motion correction job and CTF parameters were determined using Patch CTF job^20^. Only micrographs with CTF fit better than 6 Å, with astigmatism less than 1000 Å, and without significant crystalline ice contamination were considered in the next steps, effectively excluding 1091 micrographs. The accepted micrographs were filtered by Micrograph Denoiser, and the Blob Picker was used to identify particles, yielding initial 2D classes used to re-pick the whole data set with the Template Picker^20^. The false positive picks and corrupted particles were removed from the dataset by multiple rounds of reference-free 2D classification. The selected classes (449 430 particles) were subjected to initial model generation by the Ab Initio job^20^. 2D classes containing 1 845 000 particles were then subjected to Heterogeneous Refinement against 5 models generated by *ab initio* job and additional corrupted particles or particles representing alternative conformations were removed by multiple rounds of heterogeneous refinement job. This final particle set (79 525 particles) was subjected to final refinement with homogeneous refinement^20^. The reference-based motion correction was then used to improve the quality of the map, reaching a nominal resolution of 4.37 Å after homogeneous refinement according to the GSFSC0.143. The cryo-EM density map has been deposited in the Electron Microscopy Data Bank (EMDB) under accession code EMD-58943, and the fitted coordinate model in the Protein Data Bank (PDB) under accession code 32JR.

### Structure determination and structure validation

The AlphaFold model of Plg-RV was generated^21^ and used as an initial structural reference for interpretation of the cryo-EM density. A rigid-body fitting approach was carried out in UCSF Chimera X^22^. The AlphaFold model was first segmented into individual structural domains, which were subsequently treated as independent rigid bodies. Subsequently, the domains were reconnected through flexible linker regions, allowing improved correspondence between the model and the map.

The resulting model thus represents a density-guided arrangement of domains rather than a fully refined atomic structure. The final fitted model was evaluated using MolProbity^23^ (**Table S3**) to assess overall stereochemical quality. Given the moderate resolution of the cryo-EM reconstruction and the absence of full atomic refinement, the validation metrics were interpreted in the context of a rigid-body-fitted model intended primarily for visualisation of the global domain organisation. Elevated steric clashes identified during validation are consistent with the limited local resolution and the decision not to perform refinement non-supported by the experimental data.

### HDX-MS data collection and analysis

HDX-MS experiments were performed using a timsTOF SCP mass spectrometer (Bruker Daltonics) coupled to a 1290 Infinity II LC system (Agilent Technologies) with an HDX autosampler (LEAP Trajan). Deuterium labeling was initiated by 20-fold dilution of samples into D2O PBS buffer (10 mM Na2HPO4, 1.8 mM KH2PO4, 2.7 mM KCl, 137 mM NaCl, pHread 7.0, 37°C) to a final deuterium content of 81%, at labeling times of 0, 20, and 600 s. Non-deuterated controls were prepared using the corresponding H2O buffer under identical conditions. Holo-state samples were prepared by incubation of Plg variants with SAK at a 1:2 molar ratio (Plg:SAK) for 1 h at 37°C prior to deuterium labeling, matching the conditions used in the Apo-state HDX analyses. Labeling reactions were quenched by addition of an equal volume of quench buffer (3 M guanidine HCl, 250 mM TCEP, 0.5 M glycine, pH 2.3). Peptic digestion was performed online at 10°C using an immobilized dual-protease column composed of Aspergillus niger prolyl endoprotease (AnPep) and pepsin in a 1:2 ratio (Affipro). Resulting peptides were trapped and desalted online on a peptide microtrap (Phenomenex UHPLC fully porous polar C18, 2.1 mm) for 3 min, then separated on an analytical column (Phenomenex Luna Omega Polar C18, 1.6 µm, 100 × 1.0 mm, 100 Å) maintained at 2°C.

LC-MS data were processed using Data Analysis v6.1 (Bruker Daltonics) for peak picking and feature extraction. Peptide identification was performed by searching tandem mass spectra using Mascot against a database containing the sequences of all Plg variants and SAK, supplemented with common contaminants (cRAP, http://ftp.thegpm.org/fasta/cRAP). Search parameters were: precursor ion mass tolerance 10 ppm, fragment ion mass tolerance 0.05 Da, no enzyme specificity, maximum two missed cleavages, no fixed or variable modifications. Peptide-level false discovery rate was set to 1%.

Deuterium uptake at the peptide level was extracted and quantified using DeutEx (Bruker Daltonics). Back-exchange correction and residue-level relative fractional uptake (RFU) were computed from overlapping peptides using PyHDX v0.4.3^24^, with full-deuterium controls as the upper reference state and non-deuterated controls as the lower reference state.

Plg-CAT was excluded from HDX-MS analysis. As described in the Biochemical characterization section, Plg-CAT preparations consistently contained plasmin-derived proteolytic fragments arising from production-phase nicking, rendering the preparation heterogeneous and residue-level HDX-MS interpretation unreliable.

The mass spectrometry HDX data have been deposited to the ProteomeXchange (PX) Consortium^25^ via Proteomics Identifications (PRIDE)^26^ partner repository with the dataset identifier PXD080828. For peer review, the dataset can be accessed through the PRIDE repository using the following credentials: Project accession: PXD080828; Reviewer access token: 8P6lTbd5LTbY

## Results

### Production of full-length Plg variants in mammalian cells

We set up recombinant production and purification of full-length Plg variants (Plg-RV and Plg-CAT) expressed in suspension-adapted HEK293 EBNA cells^27^ (**Fig. 1A**). Despite extensive optimization, we were unable to obtain detectable expression of Plg-WT under any tested conditions, including attempts to suppress Plm activity using ε-aminocaproic acid (EACA)^28^ or aprotinin^29,30^.

**Fig. 1.**
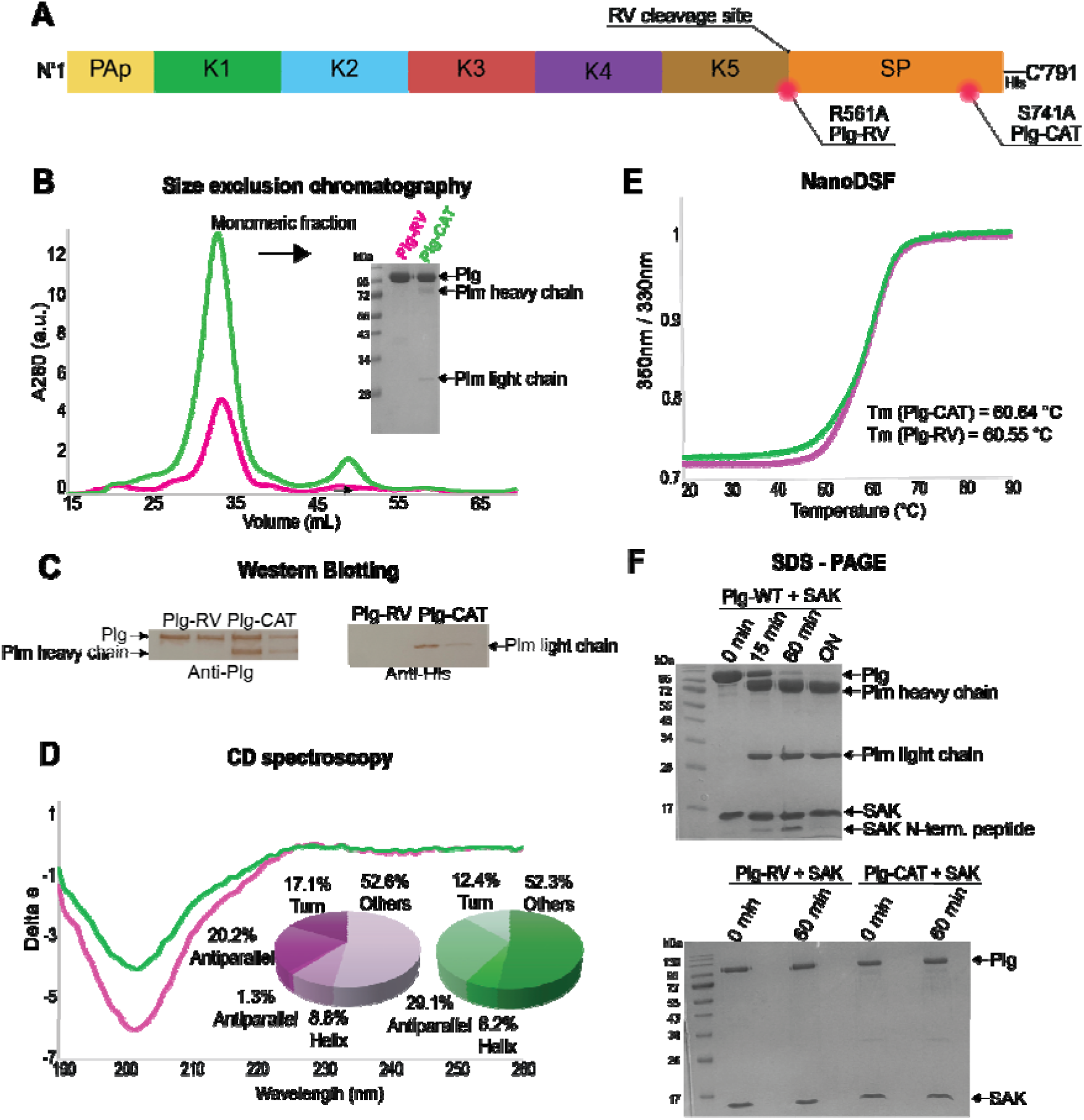
Biophysical and biochemical characterization of Plg variants. **(A)** Schematic representation of Plg topology. Individual domains are highlighted in different colours, the His-tag on the C-terminal domain is depicted. Mutations for both Plg-RV and Plg-CAT variants are highlighted. **(B)** Gel filtration profiles of Plg-RV (magenta) and Plg-CAT (green). The main peak at elution volume of ∼33 mL corresponds to the monomeric fraction of full-length Plg. Inset: SDS-PAGE analysis of the collected monomeric fractions under reducing conditions. Plg-CAT variant shows additional bands corresponding to plasmin (Plm) fragments (heavy and light chains). **(C)** Western blot characterization of purified Plg-RV and Plg-CAT. Left: Detection with anti-Plg antibody confirms the presence of full-length proteins and the N-terminal heavy chain fragment in Plg-CAT. Right: Detection with anti-His antibody identifies the C-terminal light chain fragment in Plg-CAT, consistent with the C-terminal His-tag localization **(D)** Far-UV CD spectra. The overlapping spectra indicate similar secondary structure compositions. Inset: Pie charts representing the distribution of secondary structure elements (β-sheets, α-helices, turns, and others) as estimated by the BeStSel algorithm^13^. **(E)** Thermal stability analysis showing the normalized ratio of fluorescence at 350 nm/330 nm as a function of temperature. Both variants exhibit nearly identical melting temperatures (Tm≈60.6°C), indicating correct folding and comparable global stability. **(F)** Time-course Plg activation by SAK. SDS-PAGE gels showing the incubation of Plg variants with SAK. Plg-WT (commercially sourced) is converted to plasmin within 60 min. In contrast, both recombinant variants Plg-RV and Plg-CAT remain intact, showing no proteolytic processing even after 60 min incubation. All experiments were performed in 20 mM TRIS and 20 mM NaCl, pH = 8.

Both Plg-RV and Plg-CAT recombinant variants were purified from culture supernatant using affinity chromatography followed by gel filtration (**Fig. 1B**). This protocol yielded a monodisperse, monomeric fraction for Plg-RV. In contrast, Plg-CAT consistently contained additional proteolytic fragments corresponding to Plm-derived heavy and light chains, as confirmed by SDS-PAGE, corresponding to the C-terminal light chain (∼25.2 kDa) and the N-terminal heavy chain (∼63 kDa) (**Fig. 1B**). Because the S741A substitution renders the protease domain of Plg-CAT catalytically inactive, the variant cannot undergo autocatalytic activation; the two-chain fragments therefore reflect limited nicking of the intact activation loop by an exogenous protease, most plausibly a low level of host-cell or medium-derived proteolytic activity acting over the several days of cultivation and purification. Consistent with this, Plg-RV (R561A), in which the scissile Arg561-Val562 bond is abolished, remained fully intact, indicating that the observed cleavage is specific to the activation loop rather than non-specific degradation.

Using an anti-Plg antibody, we detected both full-length Plg-RV and Plg-CAT and the N-terminal heavy chain of Plg-CAT. An anti-His antibody detected the C-terminal light chain of Plg-CAT, consistent with the localization of the His-tag on the C-terminus of the protein. The absence of such fragments in Plg-RV indicates that a proteolytic cleavage into Plm chains occurs in Plg-CAT but not in Plg-RV (**Fig. 1C**).

### Biophysical quality control

Biophysical characterization confirmed that both Plg-RV and Plg-CAT are correctly folded and structurally intact. Far-UV CD spectra indicate that both proteins are predominantly composed of β-sheets and disordered regions, which is characteristic of the multi-kringle domain architecture of plasminogen^5^. For Plg-RV, the analysis revealed a total antiparallel β-sheet content of 21.4% (consisting of 20.2% right-twisted and 1.3% relaxed strands) and a relatively low α-helical content of 8.8% (all identified as distorted helices). The remaining structure consists of turns (17.1%) and "others" (52.6%), the latter reflecting the flexible linker regions between the individual domains. Plg-CAT exhibited a similar structural profile, albeit with a higher proportion of antiparallel β-sheets (29.1%, entirely right-twisted) and a marginal decrease in helical content (6.2%). The disordered components for Plg-CAT were estimated at 12.4% for turns and 52.3% for "others" (**Fig. 1D)**. Despite the presence of minor proteolytic fragments in the Plg-CAT preparation, the CD spectra and the resulting secondary structure distribution for both variants are highly consistent with each other. NanoDSF analysis revealed nearly identical thermal stability profiles (T_m_ Plg-RV = 60.55°C, T_m_ Plg-CAT = 60.64°C) (**Fig. 1E**). Together, these results demonstrate successful production of structurally stable plasminogen variants, while also revealing intrinsic differences in proteolytic susceptibility between Plg-RV and Plg-CAT.

### Revealing differential susceptibility to SAK-mediated activation

To evaluate whether the differences in produced Plg variants translate into altered functional behavior in the presence of SAK, we next investigated Plg activation and interaction with SAK. Incubation of commercially sourced Plg-WT with SAK (in molar ratio 1 Plg : 2 SAK) resulted in a rapid N-terminal cleavage of SAK and efficient conversion of Plg into Plm. According to time-course experiments, the conversion was complete within an hour.

In contrast, both recombinant variants exhibited markedly altered responses. Plg-RV remained completely resistant to proteolytic conversion, consistent with its design as a cleavage-resistant variant at the activation site (R561A). Unexpectedly, Plg-CAT also remained intact under identical conditions (**Fig. 1F**). These differences suggest that Plg variants differ in their susceptibility to SAK-mediated activation under identical experimental conditions. To assess whether these functional differences correlate with differences in global protein conformation, we next analysed Plg-RV, Plg-CAT, and commercially available Plg-WT in solution using SAXS.

### Distinct conformational states revealed by SAXS

To assess the global conformation of Plg variants, both in-house recombinantly produced and commercially available (Athens; cat. no. 16-16-161200), we performed SAXS analyses under the same near-physiological PBS buffer (10mM Na_2_HPO_4_, 1.8 mM KH_2_PO_4_, 2.7 mM KCl, 137 mM NaCl, pH 7.4). All datasets showed a stable scattering profile without signs of aggregation. However, substantial differences were already evident in the low-q region, where Plg-RV and Plg-CAT exhibit a steeper decay of intensity compared to the more gradually decreasing profile of Plg-WT (**Fig. 2A**). Kratky plot analysis revealed a pronounced bell-shaped profile for both recombinant variants (Plg-RV and Plg-CAT), indicative of compact and well-folded particles. On the other hand, Plg-WT lacked a defined maximum, consistent with an extended or highly flexible conformation^31^ (**Fig. 2B**).

**Fig. 2.**
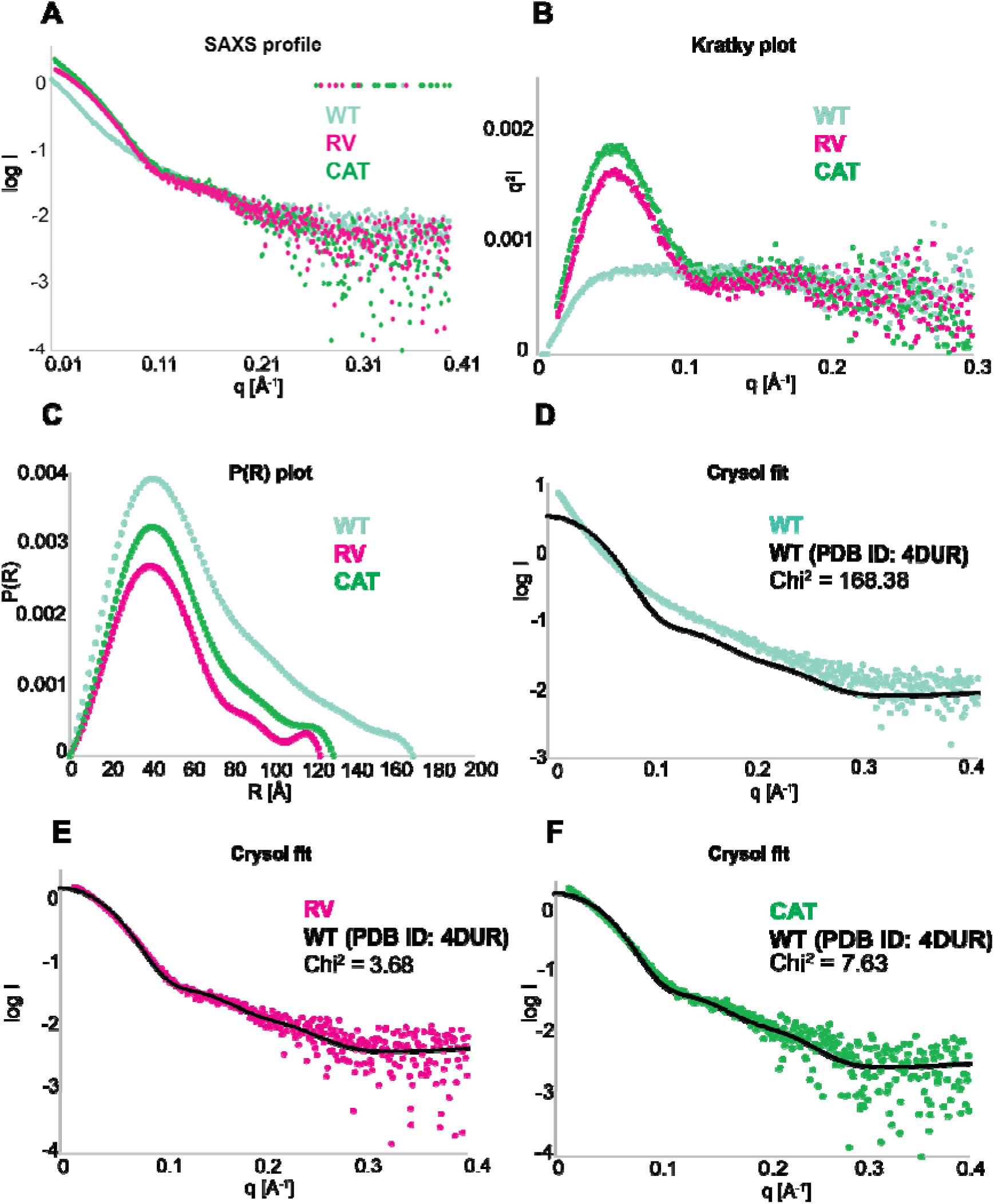
SAXS characterization of Plg variant conformations in solution. **(A)** SAXS scattering profiles for Plg-RV (magenta), Plg-CAT (green), and commercial Plg-WT (light blue). The distinct slopes in the low-q region indicate significant differences in the overall particle dimensions between the recombinant variants and the Plg-WT protein. **(B)** Kratky plots used to assess molecular flexibility and compactness. The bell-shaped peaks for Plg-RV and Plg-CAT are characteristic of well-folded, compact globular proteins. In contrast, the plateau and lack of a distinct maximum for Plg-WT suggest a more extended or flexible conformation. **(C)** Pair distance distribution functions, P(r), derived from the scattering data. The maximum particle dimension (*D*_max_) for Plg-RV and Plg-CAT is approximately 120-130 Å, while Plg-WT exhibits a larger *D*_max_ of ∼170 Å, consistent with an open, elongated structure. **(D)** CRYSOL fit comparing experimental SAXS data (dots) with theoretical scattering (black line) calculated from the crystal structure of closed plasminogen (PDB ID: 4DUR). Plg-WT shows a very poor fit (χ2=168.4), confirming it does not adopt the closed conformation. **(E)** Plg-RV shows a high degree of agreement with the closed model (χ2=3.7). (F) Plg-CAT exhibits a reasonable fit (χ2=7.6), where the slightly higher χ2 compared to RV likely reflects the presence of minor plasmin fragments previously identified. All measurements were performed in PBS (pH 7.4) at 20 °C.

Pair distance distribution functions further supported these observations, with Plg-RV and Plg-CAT exhibiting *D*_max_ values of 120-130 Å, whereas the Plg-WT showed a substantially larger *D*_max_ value of ∼170 Å (**Fig. 2C**). *Ab initio* shape reconstruction by DAMMIN^17^ yielded compact and consistent molecular envelopes for both recombinant variants, whilst the Plg-WT displayed an extended and less defined shape (**Fig. S3**). As shown in **Fig. 2D-F**, the crystal structure of Plg well agreed with the Plg-RV data (χ^2^ = 3.7), worse agreement for Plg-CAT (χ^2^ = 7.6), and a very poor fit for Plg-WT (χ^2^ = 168.4).

Together, these data demonstrate that recombinant variants (Plg-RV and Plg-CAT) predominantly adopt a compact closed conformation in solution, whereas the commercially available plasma-serum-purified counterpart populates an open-like state under identical conditions. These results could explain the poor behavior of commercial plasma-derived Plg in our initial structural experiments. In terms of conformational homogeneity, the Plg-RV variant appeared suitable for further structural investigation.

### Cryo-EM structure of Plg-RV

Next, we explored the suitability of Plg-RV for cryo EM single-particle analysis. We mixed Plg-RV with a molar excess of SAK, with an emphasis on capturing the Plg-RV-SAK complex. The particles displayed good distribution, without the signs of aggregation or preferred orientation (**Fig. 3A**). Cryo EM data were collected on a Titan Krios microscope and processed in cryoSPARC using a standard single particle analysis pipeline and the final cryo-EM map was refined to an overall resolution of 4.37 Å (FSC0.143)^32^.

**Fig. 3.**
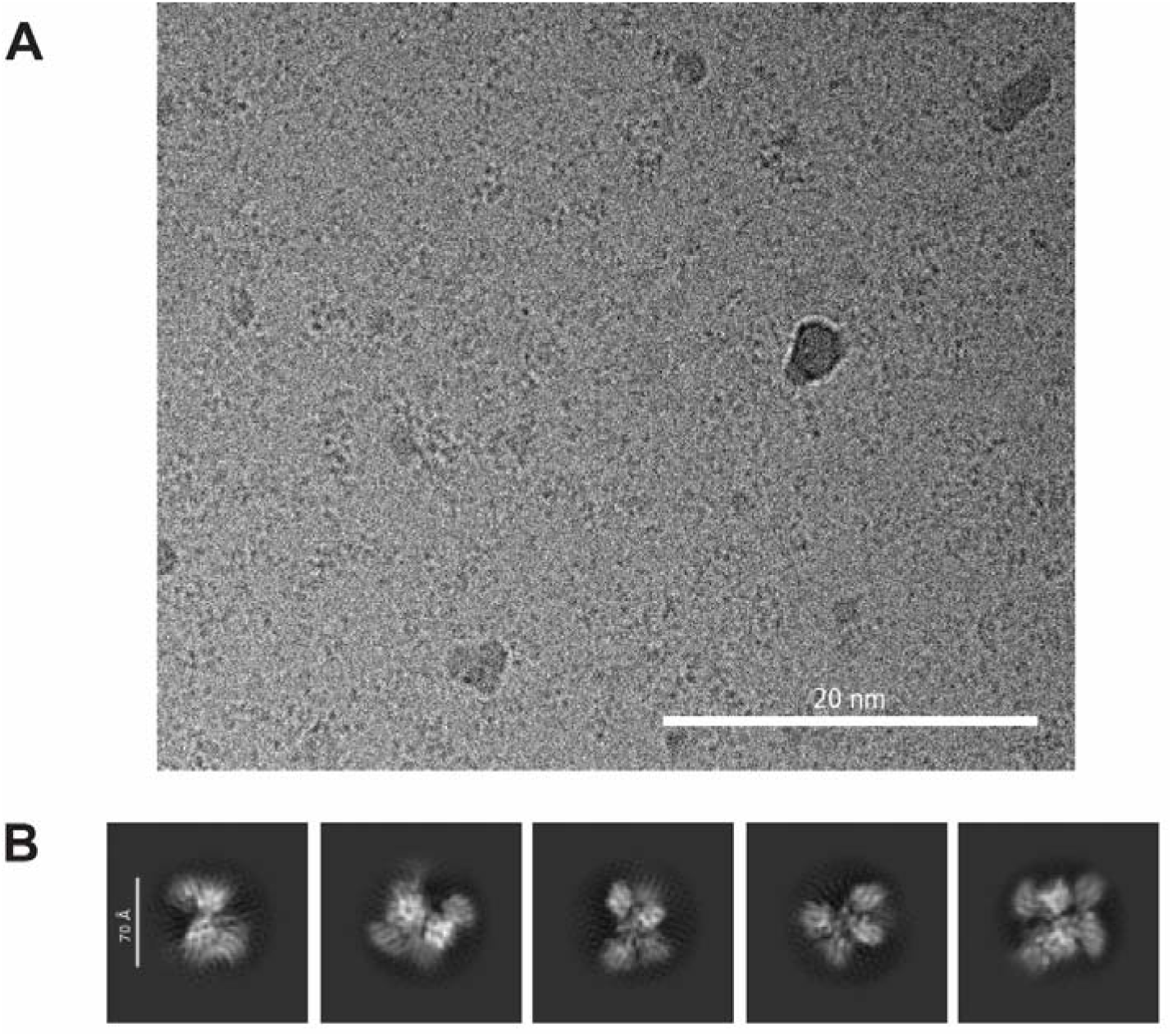
Cryo-EM data quality and 2D classification. **(A)** Representative cryo-EM micrograph of vitrified sample on a Quantifoil R1.2/1.3 grid. Particles are evenly distributed across the vitreous ice layer. The scale bar represents 200 Å. **(B)** Selected 2D classes show diverse viewing angles and well-resolved internal features, consistent with the compact, globular architecture of the closed plasminogen conformation. The scale bar represents 70 Å. These classes were used as input for *ab initio* 3D reconstruction.

The resulting density map revealed a compact, closed architecture of Plg-RV, lacking the SAK partner (**Fig. 4A,B**). The overall architecture of Plg-RV is consistent with the autoinhibited conformation previously described by X-ray crystallography^5^. The overall shape and domain arrangement are in good agreement with the SAXS-derived envelope (**Fig. S4**), providing independent validation of the closed conformation observed in solution. The map clearly defines the global organisation of the molecule, with well-resolved core regions and progressively lower local resolution towards peripheral domains and flexible linker regions.

**Fig. 4.**
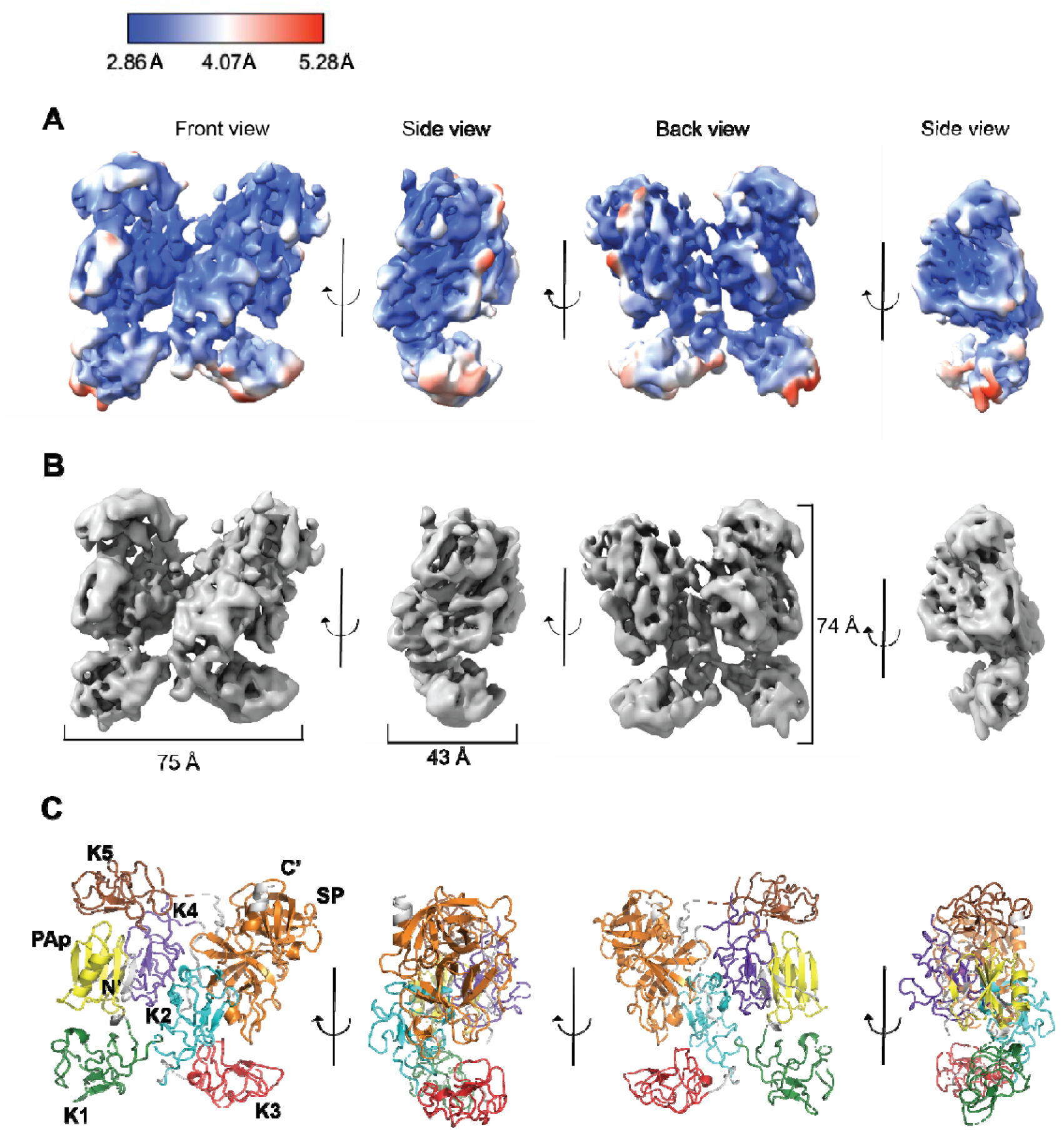
Cryo-EM structure of Plg-RV. **(A)** Cryo-EM density map of Plg-RV shown in multiple orientations, coloured according to local resolution as estimated in CryoSPARC. **(B)** Corresponding views of the cryo-EM density map displayed at a uniform threshold. Approximate dimensions of the particle are indicated. **(C)** Rigid-body fitted model of Plg-RV. Individual domains coloured and labelled, including the Pan-apple (PAp) domain, kringle domains (K1–K5), and the serine protease (SP) domain. EMDB ID: EMD-58943 and PDB ID: 32JR.

The Plg-RV model was used as a starting structural model and fitted into the cryo-EM density. Before the fitting, the model was segmented into individual domains, which were adjusted independently to achieve optimal agreement with the experimental density, to fit the Plg-RV domain arrangement recognised in the cryo-EM map. The resulting model adopts a compact arrangement consistent with the closed, autoinhibited conformation of Plg (**Fig. 4C**). While the backbone trace is well supported by the density, several loop regions and inter-domain linkers remain poorly resolved, reflecting the intrinsic flexibility of the molecule. The model displays overall geometry consistent with a rigid-body-fitted representation at moderate resolution, as assessed by MolProbity validation^23^.

Although the nominal global resolution of the reconstruction, as estimated by gold-standard FSC^32^ in CryoSPARC^20^, reaches 4.37 Å, the practical resolution varies substantially across the map, with markedly lower local resolution in peripheral regions. This indicates substantial local variability and flexibility, particularly in peripheral regions. Overall, the cryo-EM map provides a coherent and continuous representation of the full-length Plg-RV (PDB ID: 32JR), capturing its native domain organization in solution. To the best of our knowledge, this represents the first experimental visualization of the full-length Plg architecture determined by cryo-EM.

### Comparison between cryo-EM and X-ray structures

To place the cryo-EM model of Plg-RV into structural context, we compared it with the crystal structure of full-length Plg-WT in the closed conformation (PDB ID: 4DUR^5^). Structural superposition of the two models yielded an RMSD of 2.6 Å, indicating strong overall agreement in the global fold and domain organisation (**Fig. 5**). Given the moderate local resolution of the cryo-EM reconstruction and the rigid-body fitting approach used for model interpretation, the comparison was focused primarily on the overall backbone organisation and relative positioning of major secondary structure elements rather than precise local structural features. Most β-strands and α-helices occupy equivalent positions in both models, supporting the conclusion that Plg-RV retains the overall structural arrangement previously described for Plg-WT^5^.

**Fig. 5.**
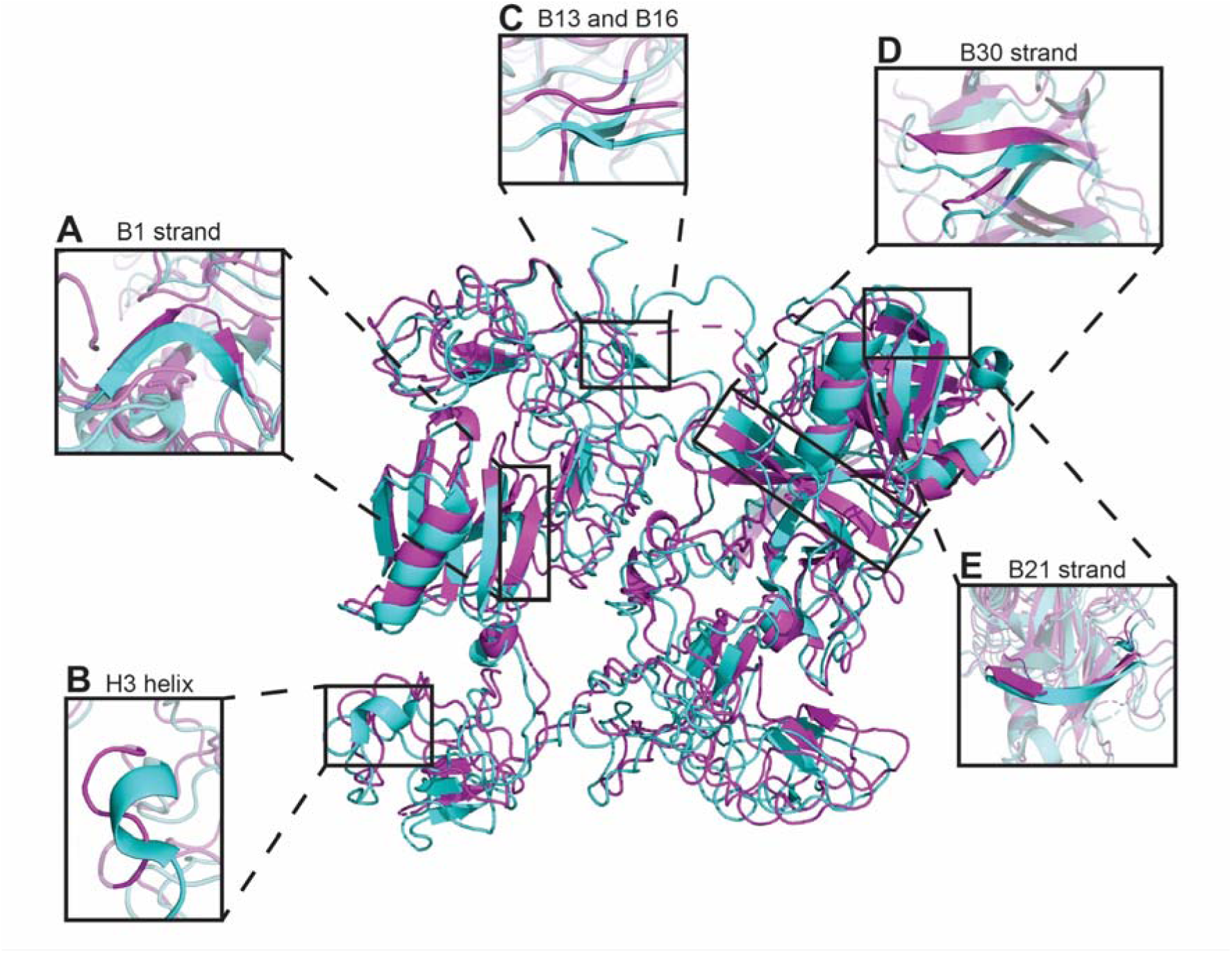
Structural comparison between Plg-RV and Plg-WT. The Plg-RV structure determined by cryo-EM in this study (PDB ID: 32JR) and Plg-WT determined by X-ray diffraction (PDB ID: 4DUR). Ribbon representation of the global structural alignment of Plg-RV (teal) and Plg-WT in the closed conformation (magenta), yielding an RMSD of 2.6 Å. Insets highlight representative regions showing local differences in secondary structure assignment or density interpretation between the models. Boxes on the central structure indicate the approximate locations of the highlighted regions. **(A)** Apparent splitting of β-strand B1 into two shorter adjacent elements (B1/B1′) in the Plg-RV cryo-EM model. **(B)** Helix H3, resolved in the crystallographic structure, is not clearly resolved in the cryo-EM model. **(C)** Reduced definition of the B13 and B16 regions in the cryo-EM model relative to the crystallographic structure. **(D)** Apparent extension of strand B30 in the Plg-RV model. **(E)** Apparent splitting of β-strand B21 into two shorter adjacent elements (B21/B21′).

Several local differences in secondary-structure assignment and density interpretation can nevertheless be observed between the cryo-EM model and the crystal structure (**Fig. 5**). These include apparent shortening or splitting of selected β-strands, reduced definition of several peripheral secondary structure elements, and local variations in loop conformations. Similar differences are also observed within parts of the serine protease domain, where several strands appear marginally extended or less continuously resolved in the cryo-EM model. Because these regions correspond to areas of lower local resolution, the observed differences likely reflect a combination of conformational heterogeneity, intrinsic flexibility, and the limitations of interpreting moderate-resolution cryo-EM density^9,10^.

Importantly, despite these local differences, the overall architecture of Plg-RV remains highly consistent with the crystallographic closed conformation of Plg-WT. The observed deviations are localised primarily in flexible peripheral regions and do not alter the global arrangement of the Pan-apple, kringle, and serine protease domains^5,6^. Together, these results support the conclusion that recombinant Plg-RV adopts the compact closed conformation.

### HDX-MS maps local conformational dynamics

To complement the global conformational characterisation provided by SAXS and cryo-EM, we performed HDX-MS on Plg-WT and Plg-RV in the apo state and following SAK incubation at Plg:SAK molar ratio of 1:2, 1 h at 37 °C. Peptide maps covering 90% and 93% of the full-length amino acid sequence were obtained for Plg-WT and Plg-RV, respectively (**Fig. 6A**). Residue-level relative fractional uptake (RFU) values were inferred from overlapping peptides using PyHDX v0.4.3^24^, see Supplementary Source HDX-MS Data file.

**Fig. 6.**
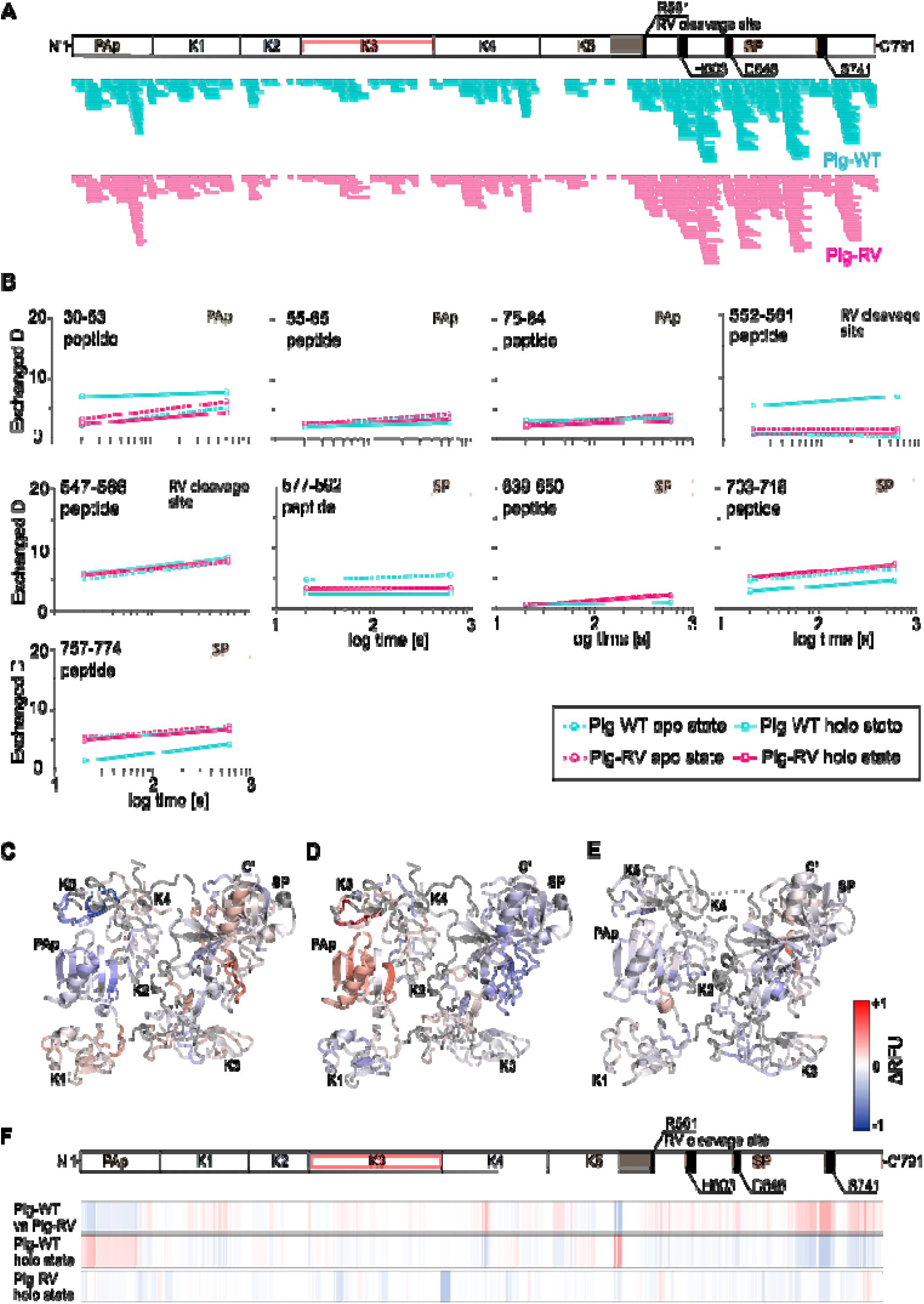
HDX-MS analysis of conformational dynamics and SAK interaction in human Plg variants. **(A)** Peptide coverage maps for Plg-WT (cyan) and Plg-RV (magenta) showing sequence coverage across the full-length protein (residues 1-791). Domain boundaries, activation cleavage site (R561, activation loop), and catalytic triad residues (H603, D646, S741) are indicated above. **(B)** Representative deuterium uptake curves for selected peptides spanning the PAp domain, activation loop region, and serine protease (SP) domain. Circles denote apo-state measurements; squares denote holo-state (SAK-bound) measurements. Plg-WT apo (cyan circle/dashed), Plg-WT holo (cyan square/solid), Plg-RV apo (magenta circle/dashed), Plg-RV holo (magenta square/solid). Domain assignment and peptide position are indicated for each panel. **(C-E)** Per-residue differential relative fractional uptake (ΔRFU) mapped onto the closed-conformation crystal structure of full-length human plasminogen (PDB: 4DUR). Colour scale: red, positive ΔRFU (increased deuterium uptake); blue, negative ΔRFU (decreased deuterium uptake); white, no difference. **(C)** Plg-WT minus Plg-RV in the apo state, reflecting conformational dynamics differences between the open (WT) and closed (RV) conformations. **(D)** Plg-WT holo minus Plg-WT apo. Because Plg-WT was largely converted to plasmin under these conditions, this comparison reflects the combined effects of activation and SAK– plasmin complex formation. Positive ΔRFU in the PAp domain indicates deprotection, whereas negative ΔRFU across the SP-domain indicates increased protection. **(E)** Plg-RV holo minus Plg-RV apo. Because Plg-RV cannot be cleaved at the activation site, this comparison reflects the structural response of the intact zymogen to SAK. Limited SP-domain protection was observed without detectable PAp-domain deprotection. **(F)** Full-protein ΔRFU heatmap for all three comparisons (Plg-WT vs Plg-RV apo; Plg-WT holo state; Plg-RV holo state), aligned to the domain map shown above. Colour scale as in C-E. Data represent single HDX-MS measurements per state; residue-level ΔRFU was resolved from overlapping peptides.

In the apo state, Plg-WT showed consistently elevated deuterium uptake relative to Plg-RV across the K5-SP linker and activation loop (residues 549−578), with maximum differences at residues 569−572 immediately downstream of the Arg561−Val562 cleavage site, and within the SP-domain core (residues 716−725; **Fig. 6B,C,F**). These data indicate that the open conformation is associated with reduced backbone protection in the activation-loop region, consistent with greater local conformational dynamics and/or solvent accessibility at a site directly involved in SAK-mediated activation. Because both proteins remained intact zymogens in the apo state, these differences cannot be attributed to Plm formation.

The holo-state comparison requires a different interpretation because the two Plg forms underwent different activation outcomes. Under the incubation conditions, plasma-derived Plg-WT was efficiently converted to plasmin, whereas cleavage-resistant Plg-RV remained an intact zymogen (**Fig. 1F**). HDX signatures for Plg-WT exhibited protection across the SP-domain (negative ΔRFU, residues 565–791) together with deprotection of the PAp domain (positive ΔRFU, residues 1–77; **Fig. 6D,E,F**). This profile is consistent with the combined effects of Plg activation and formation of the SAK–Plm complex. PAp-domain deprotection suggests weakening or release of interactions associated with the compact N-terminal arrangement, whereas SP-domain protection is consistent with engagement and/or stabilization of the protease domain in the SAK–Plm complex. In contrast, SAK induced only limited SP-domain protection in Plg-RV and no detectable PAp-domain deprotection. This pattern indicates a limited structural response involving the protease domain without the broader rearrangement, observed during activation of Plg-WT. Thus, Plg-RV provides a cleavage-resistant reference that responds locally to SAK but does not undergo conversion to Plm.

## Discussion

In this study, we present an integrative structural characterization of human Plg variants across multiple resolution scales, combining SAXS, cryo-EM, and HDX-MS. Our results demonstrate that the cleavage-resistant variant Plg-RV, recombinantly produced in mammalian cells, adopts a compact, closed conformation in solution and is amenable to cryo-EM analysis, establishing it as a structurally validated platform for investigation of the Plg−SAK system.

A central finding of this work is the conformational heterogeneity between recombinant and commercially sourced Plg preparations. SAXS analysis demonstrated that Plg-RV and Plg-CAT predominantly adopt the compact, closed conformation characteristic of the autoinhibited zymogen^5,7^, whilst the commercially available Plg-WT from Athens Research populates an extended, open-like state under identical solution conditions. This distinction is consistently reflected across all SAXS-derived parameters: the scattering profile, Kratky plot shape, maximum particle dimension, and goodness-of-fit to the closed crystal structure (PDB ID: 4DUR).

The conformational state of Plg governs its susceptibility to activation^6,7^ and influences its recognition by Plg activators, e.g. SAK.^33^ Yet this state is rarely assessed before comprehensive biochemical or structural experiments. Our data suggest that this oversight may contribute to inconsistencies in the literature and underscore the importance of conformational validation as a routine step before functional or structural studies.

The inability to produce Plg-WT recombinantly under any tested condition, including supplementation with EACA or aprotinin, is consistent with the hypothesis that unrestrained Plg activity is cytotoxic to mammalian cells.^34^ Two recombinant variants Plg-RV and Plg-CAT behaved differently in terms of preparation homogeneity. Plg-CAT consistently contained plasmin-derived heavy and light chain fragments. Because the S741A substitution abolishes catalytic activity, this nicking cannot be autocatalytic and instead reflects limited cleavage of the intact activation loop by an exogenous host cell- or culture medium-derived protease during prolonged production. The apparent discrepancy between this low-level nicking and the resistance of Plg-CAT to SAK in the activation assay (**Fig. 1F**) could reflect two mechanistically distinct processes: (i) SAK is not itself a protease but a cofactor that forms a stoichiometric activator complex with catalytically active plasmin^8^, because the protease domain of Plg-CAT is inactive, no functional SAK−plasmin activator complex can form, and the variant is therefore not converted on the timescale of the assay. The specific protease responsible for the production-phase nicking was not identified, given the catalytic inactivity of the variant, however, this activity is necessarily exogenous. This intrinsic instability rendered Plg-CAT unsuitable for single-particle cryo-EM. Plg-RV, by contrast, was produced as a homogeneous monomeric species and was therefore selected for comprehensive structural studies.

The cryo-EM reconstruction of Plg-RV at a nominal resolution of 4.37 Å represents, to our knowledge, the first experimental visualisation of full-length human plasminogen by single-particle cryo-EM. Structural comparison with the crystallographic structure PDB ID: 4DUR confirms that both adopt the same globally compact, autoinhibited architecture, with an alignment RMSD of 2.6 Å. Observed local deviations are largely confined to inter-domain linkers and peripheral loops and are most parsimoniously attributed to differences in local resolution, conformational flexibility, and the absence of crystal-contact stabilisation rather than to genuine structural differences. Consistent with this, the affected elements lie predominantly in lower-resolution regions of the map: helix H3, resolved in the crystal structure, is not clearly defined in the cryo-EM density (**Fig. 5B**), and within the serine protease domain strand B21 appears split into two shorter adjacent elements (**Fig. 5E)** while strand B30 is marginally extended relative to the wild-type model (**Fig. 5D**). Because these features fall in regions of limited local resolution, we interpret them as resolution-dependent differences in model interpretation rather than as defined conformational changes. We note that R561A lies within the serine protease domain close to the activation cleavage site, whether the substitution has any subtle local effect on the surrounding β-sheet framework cannot be determined at the present resolution and will require higher-resolution data.

Two independent observations bear on whether Plg-RV forms a complex with SAK. HDX-MS complemented the global SAXS and cryo-EM analyses by identifying local differences between the open-like Plg-WT and closed Plg-RV. In the apo state, increased deuterium uptake in the K5–SP linker and activation-loop region of Plg-WT indicated reduced local backbone protection, consistent with enhanced conformational dynamics and/or solvent accessibility. This may contribute to the increased susceptibility of the open preparation to activator-mediated cleavage, consistent with structural studies showing that the compact, closed conformation Plg-RV limits access to the Arg561–Val562 activation site, whereas conformational opening facilitates activation^5,7^. This interpretation should nevertheless remain cautious because the two preparations also differ in sequence, protein source, and production history. Moreover, HDX-MS cannot unambiguously distinguish changes in flexibility, solvent accessibility, and hydrogen-bond protection^11^.

The HDX-MS profiles obtained after SAK incubation reflect different biochemical endpoints. Plg-WT was efficiently converted to Plm and therefore reported the combined effects of proteolytic activation, conformational rearrangement, and SAK–Plm complex formation, whereas Plg-RV remained an intact cleavage-resistant zymogen. In Plg-WT, PAp-domain deprotection together with SP-domain protection was consistent with weakening of interactions associated with the compact N-terminal arrangement and stabilization of the protease domain in the SAK–Plm complex. In contrast, Plg-RV showed only limited SP-domain protection and no detectable PAp-domain deprotection, indicating a restricted structural response without the broader rearrangement associated with activation. This interpretation is consistent with the established mechanism in which SAK acquires Plg-activating activity through a Plm−SAK complexation^35,36^.

Together, these data support a model in which the open conformation not only facilitates cleavage at Arg561–Val562 but also enables the broader PAp-domain rearrangement associated with productive activation. Future high-resolution structural studies of the full-length Plg–SAK complex, building on the conformationally validated Plg-RV preparation described here, will be needed to define the precise sequence of these conformational events.

The favorable behavior of Plg-RV for cryo-EM analysis, including even particle distribution and sufficient angular coverage, can be attributed in large part to the conformational homogeneity of the preparation, as validated by SAXS. This highlights the value of orthogonal solution-state characterization as a practical screening step before committing to large-scale cryo-EM data collection. More broadly, this work demonstrates that single-particle cryo-EM analysis can be applied to Plg, despite its moderate molecular weight (∼88.4 kDa) and inherent conformational flexibility.

The cryo-EM structure presented here should be viewed as a first structural snapshot rather than a definitive endpoint. These findings should be interpreted within the bounds of the present approach. The cryo-EM map is of middle-to-low resolution and supports a domain-level, rigid-body-fitted model rather than an atomically refined structure. The absence of SAK from the reconstruction leaves the architecture of the Plg−SAK complex undefined. In addition, the open-like reference state was established using a single commercial, plasma-derived (Glu-plasminogen) preparation. While this is sufficient to demonstrate the conformational divergence central to this study, the individual contributions of plasma origin, the Glu-form, and purification history to this open state were not dissected here and will require future research.

## Acknowledgements

The authors acknowledge the support of the RECETOX Research Infrastructure (No. LM2023069) and CZECRIN (No. LM2023049), funded by the Ministry of Education, Youth and Sports (MEYS) of the Czech Republic, and the Ministry of Health of the Czech Republic - Conceptual development of research organization (MMCI, 00209805). This project was also supported by the European Union’s Horizon 2020 Research and Innovation Programme under grant agreement No. 857560 (CETOCOEN Excellence), and has received funding from the Horizon Europe programme under grant agreements No. 101087124 (ADDIT-CE), and No. 101136607 (CLARA). Additional funding was provided through the ESIF-MEYS Johannes Amos Comenius Programme under the CLARA project (No. CZ.02.01.01/00/23_029/0008437), co-financed by the European Union and MEYS, and Czech Science Foundation (GX25-17329X). Views and opinions expressed are however those of the author(s) only and do not necessarily reflect those of the European Union or REA. Neither the European Union nor the granting authority can be held responsible for any use that may be made of the information it contains. Computational resources were provided by the e-INFRA CZ and ELIXIR-CZ (No. 90254 and LM2023055 MEYS). We acknowledge CEITEC CF CEMCOF, CF BIC and CF NMR of CIISB, Instruct-CZ Centre, supported by MEYS CR (LM2023042) and European Regional Development Fund-Project „Innovation of Czech Infrastructure for Integrative Structural Biology” (No. CZ.02.01.01/00/23_015/0008175). LK is supported by the scholarship Brno Ph.D. Talent. We acknowledge Cryo-electron microscopy and tomography core facility CEITEC MU of CIISB, Instruct-CZ Centre, supported by MEYS CR (LM2023042) and European Regional Development Fund-Project „Innovation of Czech Infrastructure for Integrative Structural Biology” (No. CZ.02.01.01/00/23_015/0008175).

## Data availability

The cryo-EM density map of full-length plasminogen variant Plg-RV has been deposited in the Electron Microscopy Data Bank (EMDB) under accession code EMD-58943, and the corresponding fitted coordinate model in the Protein Data Bank (PDB) under accession code 32JR. The small-angle X-ray scattering data have been deposited in the Small Angle Scattering Biological Data Bank (SASBDB) under accession codes SASDZU3, SASDZ24, and SASDZ34. The hydrogen/deuterium exchange mass spectrometry data have been deposited to the ProteomeXchange Consortium via the PRIDE partner repository under the dataset identifier PXD080828.

## References

1. Plow, E. F., Herren, T., Redlitz, A., Miles, L. A. & Hoover-Plow, J. L. The cell biology of the plasminogen system. FASEB J. Off. Publ. Fed. Am. Soc. Exp. Biol. 9, 939–945 (1995).

2. Keragala, C. B. & Medcalf, R. L. Plasminogen: an enigmatic zymogen. Blood 137, 2881– 2889 (2021).

3. Peetermans, M. et al. Plasminogen activation by staphylokinase enhances local spreading of S. aureus in skin infections. BMC Microbiol. 14, 310 (2014).

4. Grella, D. K. & Castellino, F. J. Activation of human plasminogen by staphylokinase. Direct evidence that preformed plasmin is necessary for activation to occur. Blood 89, 1585–1589 (1997).

5. Law, R. H. P. et al. The X-ray Crystal Structure of Full-Length Human Plasminogen. Cell Rep. 1, 185–190 (2012).

6. Quek, A. J. et al. A High-Throughput Small-Angle X-ray Scattering Assay to Determine the Conformational Change of Plasminogen. Int. J. Mol. Sci. 24, 14258 (2023).

7. Xue, Y., Bodin, C. & Olsson, K. Crystal structure of the native plasminogen reveals an activation resistant compact conformation. J. Thromb. Haemost. 10, 1385–1396 (2012).

8. Parry, M. A. et al. The ternary microplasmin-staphylokinase-microplasmin complex is a proteinase-cofactor-substrate complex in action. Nat. Struct. Biol. 5, 917–923 (1998).

9. Herzik, M. A., Wu, M. & Lander, G. C. High-resolution structure determination of sub-100 kDa complexes using conventional cryo-EM. Nat. Commun. 10, 1032 (2019).

10. Wentinck, K., Gogou, C. & Meijer, D. H. Putting on molecular weight: Enabling cryo-EM structure determination of sub-100-kDa proteins. Curr. Res. Struct. Biol. 4, 332–337 (2022).

11. Vinciauskaite, V. & Masson, G. R. Fundamentals of HDX-MS. Essays Biochem. 67, 301–314 (2023).

12. Legrand, A. et al. Investigating the Conformational Flexibility of Staphylokinase Across Multiple Time Scales. 2026.02.03.703606 Preprint at 10.64898/2026.02.03.703606 (2026).

13. Micsonai, A. et al. Accurate secondary structure prediction and fold recognition for circular dichroism spectroscopy. Proc. Natl. Acad. Sci. U. S. A. 112, E3095–3103 (2015).

14. Franke, D. et al. ATSAS 2.8: a comprehensive data analysis suite for small-angle scattering from macromolecular solutions. J. Appl. Crystallogr. 50, 1212–1225 (2017).

15. Konarev, P. V., Volkov, V. V., Sokolova, A. V., Koch, M. H. J. & Svergun, D. I. PRIMUS: a Windows PC-based system for small-angle scattering data analysis. J. Appl. Crystallogr. 36, 1277–1282 (2003).

16. Svergun, D., Barberato, C. & Koch, M. H. J. CRYSOL – a Program to Evaluate X-ray Solution Scattering of Biological Macromolecules from Atomic Coordinates. J. Appl. Crystallogr. 28, 768–773 (1995).

17. Svergun, D. I. Restoring Low Resolution Structure of Biological Macromolecules from Solution Scattering Using Simulated Annealing. Biophys. J. 76, 2879–2886 (1999).

18. Valentini, E., Kikhney, A. G., Previtali, G., Jeffries, C. M. & Svergun, D. I. SASBDB, a repository for biological small-angle scattering data. Nucleic Acids Res. 43, D357–D363 (2015).

19. Mastronarde, D. N. Automated electron microscope tomography using robust prediction of specimen movements. J. Struct. Biol. 152, 36–51 (2005).

20. Punjani, A., Rubinstein, J. L., Fleet, D. J. & Brubaker, M. A. cryoSPARC: algorithms for rapid unsupervised cryo-EM structure determination. Nat. Methods 14, 290–296 (2017).

21. Jumper, J. et al. Highly accurate protein structure prediction with AlphaFold. Nature 596, 583–589 (2021).

22. Pettersen, E. F. et al. UCSF ChimeraX: Structure visualization for researchers, educators, and developers. Protein Sci. Publ. Protein Soc. 30, 70–82 (2021).

23. Williams, C. J. et al. MolProbity: More and better reference data for improved all-atom structure validation. Protein Sci. Publ. Protein Soc. 27, 293–315 (2018).

24. Smit, J. H. et al. Probing Universal Protein Dynamics Using Hydrogen–Deuterium Exchange Mass Spectrometry-Derived Residue-Level Gibbs Free Energy. Anal. Chem. 93, 12840–12847 (2021).

25. Deutsch, E. W. et al. The ProteomeXchange consortium at 10 years: 2023 update. Nucleic Acids Res. 51, D1539–D1548 (2023).

26. Perez-Riverol, Y. et al. The PRIDE database resources in 2022: a hub for mass spectrometry-based proteomics evidences. Nucleic Acids Res. 50, D543–D552 (2022).

27. Durocher, Y., Perret, S. & Kamen, A. High-level and high-throughput recombinant protein production by transient transfection of suspension-growing human 293-EBNA1 cells. Nucleic Acids Res. 30, e9 (2002).

28. Wu, T. P., Padmanabhan, K., Tulinsky, A. & Mulichak, A. M. The refined structure of the epsilon-aminocaproic acid complex of human plasminogen kringle 4. Biochemistry 30, 10589–10594 (1991).

29. Fritz, H. & Wunderer, G. Biochemistry and applications of aprotinin, the kallikrein inhibitor from bovine organs. Arzneimittelforschung. 33, 479–494 (1983).

30. Al-Horani, R. A. & Desai, U. R. Recent Advances on Plasmin Inhibitors for the Treatment of Fibrinolysis-Related Disorders. Med. Res. Rev. 34, 1168–1216 (2014).

31. Kikhney, A. G. & Svergun, D. I. A practical guide to small angle X-ray scattering (SAXS) of flexible and intrinsically disordered proteins. FEBS Lett. 589, 2570–2577 (2015).

32. Scheres, S. H. W. & Chen, S. Prevention of overfitting in cryo-EM structure determination. Nat. Methods 9, 853–854 (2012).

33. Sakharov, D. V., Lijnen, H. R. & Rijken, D. C. Interactions between staphylokinase, plasmin(ogen), and fibrin. Staphylokinase discriminates between free plasminogen and plasminogen bound to partially degraded fibrin. J. Biol. Chem. 271, 27912–27918 (1996).

34. Busby, S. J. et al. Expression of recombinant human plasminogen in mammalian cells is augmented by suppression of plasmin activity. J. Biol. Chem. 266, 15286–15292 (1991).

35. Nguyen, L. T. & Vogel, H. J. Staphylokinase has distinct modes of interaction with antimicrobial peptides, modulating its plasminogen-activation properties. Sci. Rep. 6, 31817 (2016).

36. Toul, M., Nikitin, D., Marek, M., Damborsky, J. & Prokop, Z. Extended Mechanism of the Plasminogen Activator Staphylokinase Revealed by Global Kinetic Analysis: 1000-fold Higher Catalytic Activity than That of Clinically Used Alteplase. ACS Catal. 12, 3807–3814 (2022).

