## Supplementary Information for "Open or closed? Exploring the conformational heterogeneity of human plasminogen across multiple resolution scales"

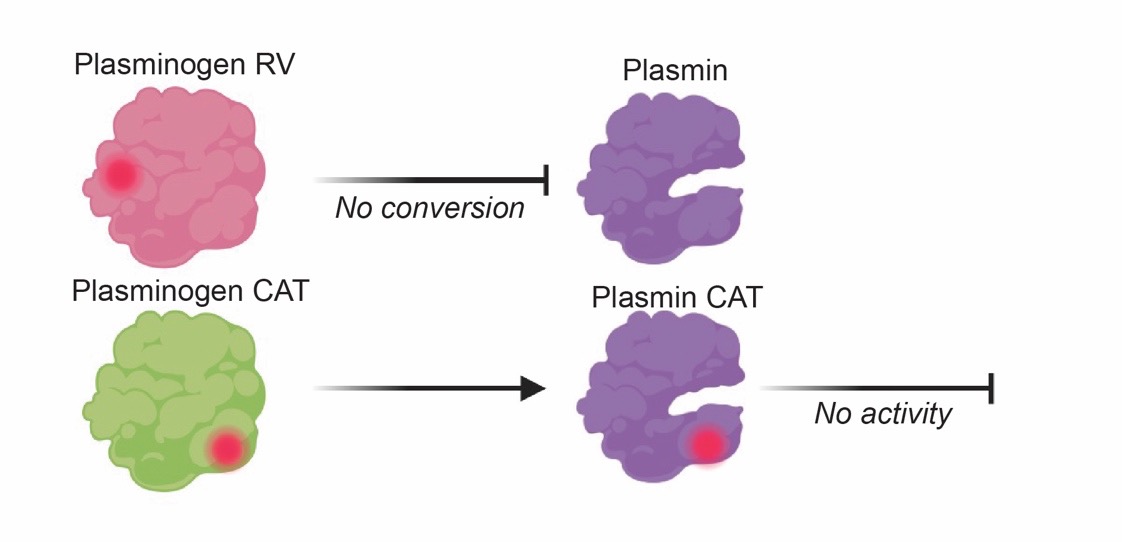

**Fig. S1.** Schematic representation of the plasminogen designed variants Plg-RV and Plg-CAT. In both variants, the selected amino acid residues were substituted with alanine to abolish their functional contribution. Plg-RV has an alanine in the “to - plasmin” cleavage site (R561A), therefore it cannot be converted to plasmin. Plg-CAT has an alanine in the active site (S741A), thus it can generate plasmin, but this plasmin would be proteolytically inactive.

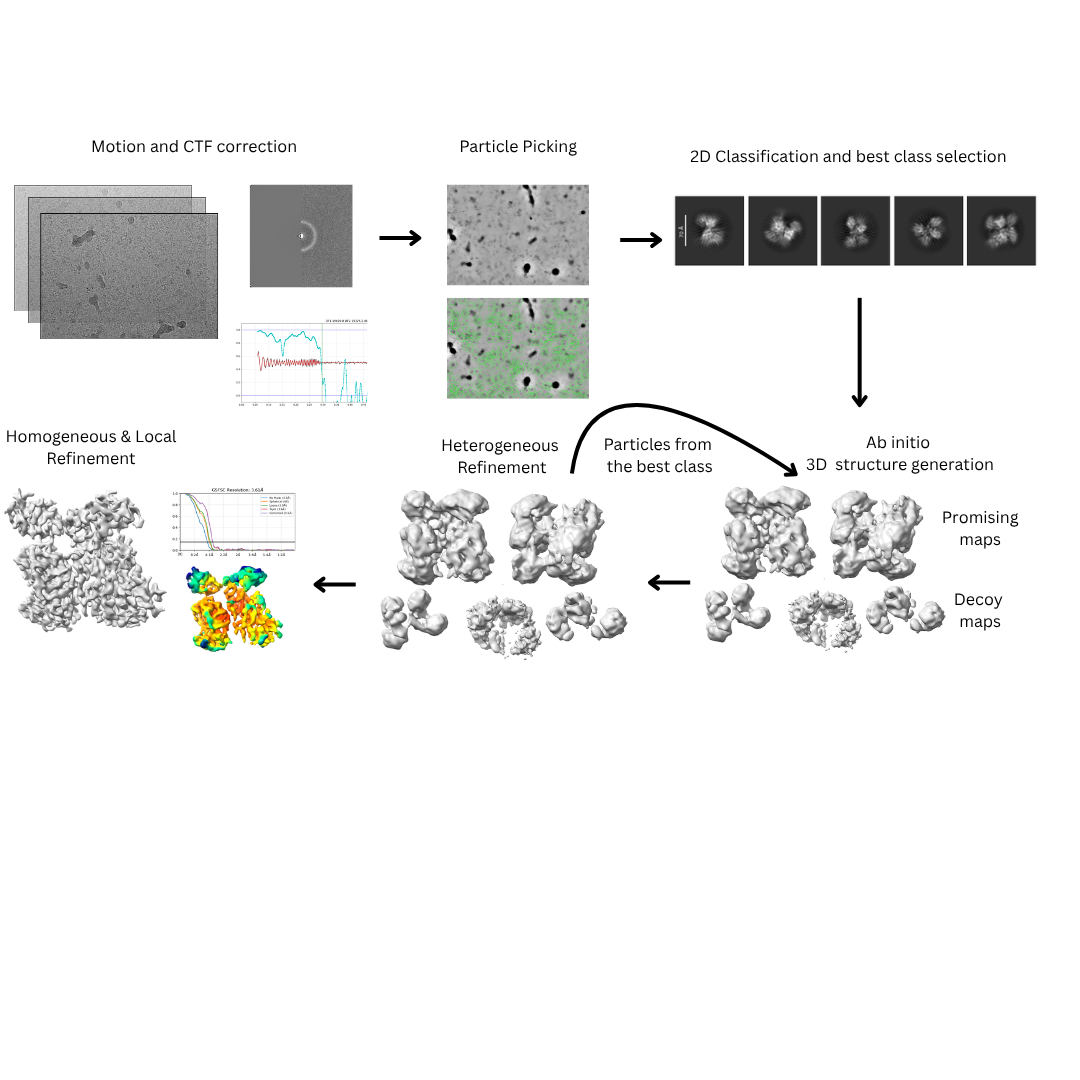

**Fig. S2.** Depiction of cryo-EM data collection and processing workflow used for structural characterization of Plg-RV variant.

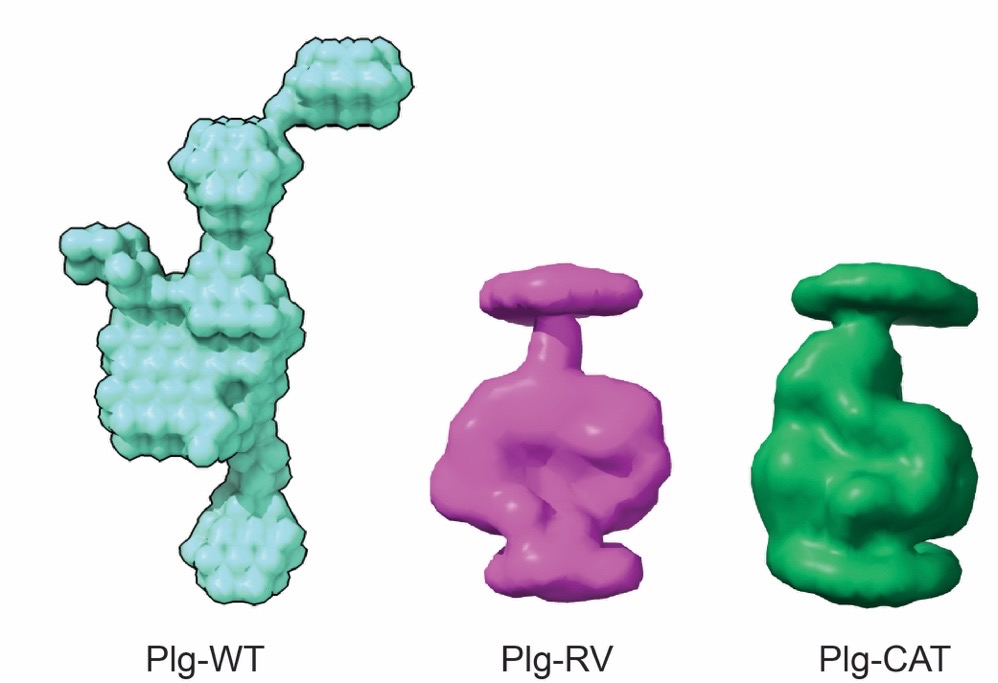

**Fig. S3.** *Ab initio* SAXS envelopes of Plg-WT (cyan), Plg-RV (magenta) and Plg-CAT (green).

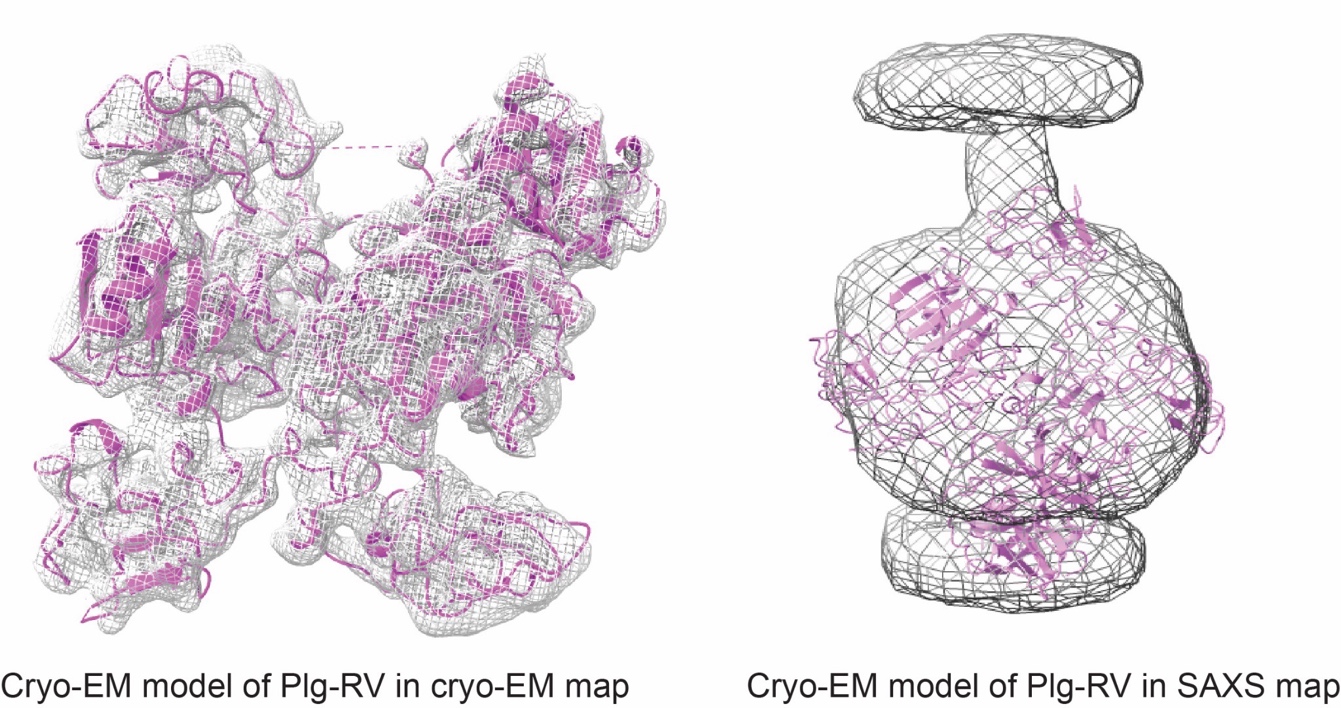

**Fig. S4.** Plg-RV model aligned in the cryo-EM map and Plg-RV model aligned in the SAXS *ab initio* molecular envelope.

**Table S1**. SAXS measurements and integral structural parameters determined by PRIMUS.

|  | **Plg-WT** | **Plg-RV** | **Plg-CAT** |
| --- | --- | --- | --- |
| Instrument | BioSAXS-2000 | BioSAXS-2000 | BioSAXS-2000 |
| Wavelength [Å] | 1.54 | 1.54 | 1.54 |
| q range [Å-1] | 0.009 - 0.65 | 0.009 - 0.65 | 0.009 - 0.65 |
| Exposure time [min] | 60 | 40 | 40 |
| Temperature [°C] | 20 | 20 | 20 |
| Concentration [mg/mL] | 2.50 | 5 | 5 |
| Rg [Å] (from Guinier) | 53.82 | 37.25 | 41.97 |
| Rg [Å] (from P(R)) | 51.37 | 37.55 | 41.02 |
| Dmax [Å] | 168 | 124 | 130 |
| Porod volume estimate [Å³] | 120 000 | 161 710 | 200 607 |
| Mw from sequence [kDa] | 88.56 | 88.36 | 88.43 |
| Primus Mw estimate [kDa] | 62.4 | 75.3 - 92.7 | 102.9 - 116 |

**Tab. S2.** Cryo-EM parameters.

| **Parameter** | **Value** |
| --- | --- |
| Microscope | Titan Krios |
| Voltage | 300 kV |
| Cs | 2.7 mm |
| Camera | K3 |
| Energy filter | Bioquantum |
| Slit width | 10 eV |
| Magnification | 165 kx |
| Pixel size | 0.5113 Å/pix |
| Dose rate | 7.5 e/Å²/s |
| Total dose | 56 e/Å² |
| Exposure time | 2 s |
| Frames | 40 |
| Defocus min | -1 |
| Defocus max | -2.4 |
| total movies | 14 330 |
| Particle picks | 5 760 000 |
| Particle after 2D | 1 845 000 |
| Particle after 3D | 79 525 |

**Table S3.** MolProbity all-atom contact and geometry validation statistics for the final PLG-RV model.

| **Category** | **Statistic** | **Value** |
| --- | --- | --- |
| All-Atom Contacts | Clashscore, all atoms | 41.89 (7th percentile, N = 1784) |
| Protein Geometry | Poor rotamers | 0 (0.00%) |
|  | Favored rotamers | 640 (98.92%) |
|  | Ramachandran outliers | 0 (0.00%) |
|  | Ramachandran favored | 695 (97.20%) |
|  | Rama distribution Z-score | 0.53 ± 0.29 |
|  | MolProbity score | 2.25 (62nd percentile, N = 27675) |
|  | Cβ deviations > 0.25 Å | 0 (0.00%) |
|  | Bad bonds | 1 / 5963 (0.02%) |
|  | Bad angles | 12 / 8094 (0.15%) |
| Peptide Omegas | Cis prolines | 5 / 58 (8.62%) |
| Low-resolution Criteria | CaBLAM outliers | 11 (1.6%) |
|  | CA geometry outliers | 10 (1.44%) |
| Additional Validations | Chiral volume outliers | 0 / 838 |
|  | Waters with clashes | 0 / 0 (0.00%) |
